# Chromatin nanoscale compaction in live cells visualized by acceptor-donor ratio corrected FRET between DNA dyes

**DOI:** 10.1101/671784

**Authors:** Simone Pelicci, Alberto Diaspro, Luca Lanzanò

**Affiliations:** Nanoscopy and Nikon Imaging Center, Istituto Italiano di Tecnologia, via Morego 30, 16163 Genoa, Italy; Department of Physics, University of Genoa, via Dodecaneso 33, 16143 Genoa, Italy

**Keywords:** FRET, FLIM, phasors, DNA dyes, nanoscale chromatin organization

## Abstract

Chromatin nanoscale architecture in live cells can be studied by Forster Resonance Energy Transfer (FRET) between fluorescently labeled chromatin components, such as histones. A higher degree of nanoscale compaction is detected as a higher FRET level, since this corresponds to a higher degree of proximity between donor and acceptor molecules. However, in such a system the stoichiometry of the donors and acceptors engaged in the FRET process is not well defined and, in principle, FRET variations could be caused by variations in the acceptor-donor ratio rather than distance. Here we show that a FRET value independent of the acceptor-donor ratio can be obtained by Fluorescence Lifetime Imaging (FLIM) detection of FRET combined with a normalization of the FRET level to a pixel-wise estimation of the acceptor-donor ratio. We use this method to study FRET between two DNA binding dyes staining the nuclei of live cells. We show that acceptor-donor ratio corrected FRET imaging reveals variations of nanoscale compaction in different chromatin environments. As an application, we monitor the rearrangement of chromatin in response to laser-induced micro-irradiation and reveal that DNA is rapidly decompacted, at the nanoscale, in response to DNA damage induction.

## 1. Introduction

A major function of chromatin is the organization and compaction of long genomic DNA within the confined space of the eukaryotic nucleus. Chromatin nanoscale architecture, namely the organization of histones into higher-order structures of varying level of compaction, has an important role in the regulation of many genomic processes. For instance, the modulation of gene activity, which involves the packaging of the genome into transcriptionally active and inactive sites, is dependent upon chromatin morphological organization [1][2]. Another example is represented by the chromatin rearrangements that occur during the DNA damage response (DDR) to facilitate access of specific proteins, implicated in various DNA repair pathways and in the maintenance of genomic stability [3][4][5]. How such structures form and behave in various cellular processes remains unclear and how genomes are structured at the nanoscale, especially in living cells, remains unknown.

Historically, different chromatin compaction states have been broadly divided into heterochromatin and euchromatin, owing to their association with the density of their appearance with optical light microscopy (OLM) or electron microscopy (EM) [6][7]. Heterochromatin was first defined as the fraction of chromatin that remains condensed after mitosis while euchromatin has been described as low density, relatively decompacted chromatin, which includes mostly active regions rich in genes and regulatory sequences [8]. The recent development of the so-called super-resolution fluorescence microscopy (SRM) techniques, including stimulated emission depletion microscopy (STED) [9], structured illumination microscopy (SIM)[10], and localization microscopy, such as photoactivated localization microscopy (PALM)[11] and stochastic optical reconstruction microscopy (STORM) [12][13], have extended the ultimate resolving power of optical microscopy far beyond the diffraction limit, facilitating access to the organization of chromatin at the nanoscale by optical means [14][15][16][17][18][19]. For instance, STORM has been used to visualize chromatin higher order organization in single cell nuclei, revealing that both nucleosomes and DNA associate in heterogeneous nanodomains [16][20], and that distinct epigenetic states have a different nanoscale chromatin architecture [15][21]. Notably, PALM has been used to visualize the dynamics of higher-order chromatin structures in live cells[17].

An alternative strategy to get information on the nanoscale chromatin environment, without the help of SRM, is based on the use of fluorescence spectroscopy techniques. For instance, techniques like fluorescence recovery after photobleaching (FRAP) and fluorescence correlation spectroscopy (FCS) have been widely used to measure molecular diffusion within the nucleus of live cells [22]. Even if limited by diffraction, these methods detect differences in the diffusion of nanometer-sized probes and can be used to indirectly infer properties of the nanoscale chromatin architecture [23][24][25][26][27]. FCS can be eventually coupled with STED to probe diffusion at sub-diffraction spatial scales [28][29]. Other examples are Fluorescence Lifetime Imaging Microscopy (FLIM) and fluorescence anisotropy imaging, which have been used to monitor alterations of viscosity in the local environment of a fluorescent probe and relate them to the chromatin condensation state [30][31]. But probably the most striking example of achieving the nanoscale through fluorescence spectroscopy is represented by Förster resonance energy transfer (FRET), a process that can occur between an excited donor and an acceptor molecule when the two fluorophores are within ∼10 nm distance [32]. Because of this property FRET is often considered a ‘spectroscopic nanoruler’ and is the method of choice for the detection of protein-protein interactions in live cells [33][34][35].

The sensitivity to nanometer distances makes FRET especially interesting in the context of the highly packed chromatin environment. A quantitative FRET approach to assay nanoscale chromatin compaction was originally developed by Lleres and coworkers [36]. This assay defines the nanoscale proximity between nucleosomes, measured through FLIM-based detection of FRET between stably incorporated GFP-H2B and mCherry-H2B histones. This FRET assay has been used to reveal distinct domains and quantitatively discriminate different levels of nanoscale chromatin compaction in live HeLa cells [36] and in living C. Elegans as a model system [37]. More recently, the same FRET assay has been applied to measure chromatin organization in live cells in combination with the phasor analysis of FLIM[38][39]. Coupling this technology with laser micro-irradiation allowed to identify the DDR-dependent chromatin architectural changes that occur in response to DNA double-strand breaks (DSBs) [39]. All these works clearly show that FRET can be a powerful tool to map nanoscale chromatin compaction in vivo.

Nevertheless, there are some limitations in the reported chromatin compaction FRET assays. First, the use of fluorescent proteins implies in general a longer time to prepare samples, required for inducing a transient expression of the proteins, or the establishment of a stable cell line. In addition, the induction of transient or stable fluorescent protein expression might be challenging in some specific cell lines, limiting the applicability of the method. A second important consideration is that, in the chromatin FRET assay, the effective number of donors and acceptors involved in the FRET interaction is not well defined. This is quite different from the FRET detection of protein-protein interactions, in which the stoichiometry of the putative protein clusters is often predictable, at least to some extent. In particular, in the case of fluorescent histones, up to 20-fold variations in the relative acceptor-donor expression level were recently reported [40]. A high variability in the number of donor and acceptor molecules may generate by itself additional variations of FRET not necessarily linked to variations in the average donor-acceptor distance [41][42][43][44][45]. If this is the case, the measured FRET level should be corrected for the relative acceptor donor abundance.

Here, we introduce a novel FRET assay that provides a FRET value independent of the acceptor-donor ratio. The assay is based on the FRET between two DNA-binding dyes, namely Hoechst 33342 (donor) and Syto 13 (acceptor), rather than between fluorescently labeled histones. In order to provide an accurate FRET level, we monitor variations of the lifetime of the donor by frequency-domain FLIM, and normalize the FRET efficiency to the relative acceptor-to-donor abundance. We show that, thanks to this correction, the method provides consistent spatial maps of nanoscale chromatin compaction independently of the local donor and acceptor concentrations. We validate the method by quantification of different degrees of chromatin compaction in live interphase nuclei, distinguishing different density patterns, both in physiological and hyperosmolar environment. As an application, we study changes in nanoscale chromatin architecture during the DNA damage response (DDR), generated by stimulation with laser UV-microirradiation, inside nuclear defined regions.

## 2. Materials and methods

### 2.1. Cell Culture and Treatments

HeLa cells were cultured in a flask in Dulbecco’s modified Eagle’s medium (DMEM) supplemented with 10% FBS, 2 mM L-glutamine and 1% penicillin/streptomycin in a humidified incubator at 37°C with 5% CO2. Subsequently, cells were plated on a Ibidì µ-slide 8-well chamber and let grow overnight. Cells were washed in Phosphate Buffer Saline (PBS 1X, pH 7.4; Thermofisher Scientific) and stained with 2µM Hoechst 33342 (Thermofisher Scientific) (donor only sample) or with 2µM Hoechst 33342 and 2µM Syto 13 (Thermofisher Scientific) (donor-acceptor sample) and left incubating for 25 min at 37 C. For FRET measurements, cells were observed without any washing step, i.e., leaving the fluorophore diluted in DMEM.

For hyperosmolar experiment, HeLa cells were washed in PBS 1X and hyper-compacted chromatin (HCC) formation was induced by incubating HeLa cells in a hyper-osmolar medium at osmolarities ∼570 mOsm for 25min at 37°C with 5% CO2. Afterwards cells were stained with 2µM Hoechst 33342 (donor only) or with 2µM Hoechst 33342 and and 2µM Syto 13 (donor-acceptor sample) and left incubating for 20 min at 37°C in hyperosmolar solution. As a standard protocol, 1 ml 20 X PBS (2.8 M NaCl, 54 mM KCl, 130 mM Na2HPO4, 30 mM KH2PO4 in H2O, pH adjusted with HCl to 7.4) was diluted with 19 ml standard culturing medium (290 mOsm) to yield an osmolarity of 570 mOsm [46].

For monitoring the DNA damage response, cells were transiently transfected with PARP1 chromobody tagged-RFP (Chromotek), according to QIAGEN Effectene protocol and imaged 24 h after transfection.

### 2.2. Microscopy

FLIM-FRET data were acquired with Nikon’s A1R MP confocal and multiphoton microscope, coupled to an ISS A320 frequency-domain FastFLIM box to acquire the lifetime data. A Nikon Plan Apo VC 100X Oil DIC N2 objective, NA 1.45, was used for all the measurements. The donor fluorophore was excited at 405 nm. This wavelength caused also direct excitation of the acceptor. The fluorescence signal was split between two hybrid photodetectors, with the following emission band-pass filters in front of each: 450/50 (Hoechst 33342) and 585/40 (Syto 13), respectively. We simultaneously acquired intensity and lifetime data by scanning with a 80 MHz pulsed laser beam (405 nm, PDL 800-D, PicoQuant). The frame size was set to 512×512 pixels, with a pixel size of ∼0.05 μm. The scanning pixel-dwell time was set at 12.1 μs/pixel. Each FLIM image was obtained as the average of 20 frames corresponding to an acquisition time of ∼1 min.

The FLIM data acquisition was managed by the ISS VistaVision software. In frequency domain FLIM, the lifetime is determined from the phase delay and the de-modulation of the fluorescence emission with respect to a modulated excitation signal [47]. For each pixel, the FLIM system records a value of phase (φ) and modulation (M) at multiple frequencies with respect to the excitation signal. All the data were analyzed at the frequency of 80 MHz. The raw FLIM data were visualized in the phasor plot where *g*=*M* cos(φ) and *s*=*M* sin(φ) [38]. Calibration of the system was performed by measuring Fluorescein in 1M NaOH (pH 9.0), which has a known single exponential lifetime of 4.04 ns. Before each experiment, we calibrated the donor channel using a solution of Alexa Fluor 405 (Thermofisher) which is excited at the same excitation wavelength of the donor (Hoechst 33342). We determined that Alexa Fluor 405 in DMSO has a single exponential lifetime of 3.5 ns (Supplementary Fig.4). For each measurement, the following four images were exported for further processing on ImageJ [48]: the intensity in the donor channel *I*_1_(*x,y*), the intensity in the acceptor channel *I*_2_(*x,y*), the phasor coordinate *g*(*x,y*) in the donor channel and phasor coordinates *s*(*x,y*) in the donor channel.

### 2.3. Laser micro-irradiation

For induction of DNA damage by laser micro-irradiation, we used the 405 nm-laser beam of the Nikon’s A1R MP confocal and multiphoton microscope. The laser power was set at 80% and the laser beam was focused on a selected region of interest (ROI) of the nucleus (15 µm × 6 µm) for a total micro-irradiation time of 40 s. For monitoring the response of PARP-1 to DNA damage induction, a 65 s time-lapse movie was recorded (256 × 256 pixels, 9.5 µs/pixel, 634 frames). FLIM-FRET microscopy was performed in parallel using the microscope and acquisition settings described above. FLIM-FRET acquisitions were recorded immediately after laser micro-irradiation.

For cells analyzed by immunostaining, the induction of DNA damage was set with a laser power at 100%, on a selected ROI of size 3 µm × 3 µm, for a total micro-irradiation time of 20 s. Micro-irradiated cells were fixed within ∼5 min after micro-irradiation.

### 2.4. Cell fixation and immunostaining

Cells were fixed with 4% formaldehyde in PBS 1× for 15 min and washed several times with PBS 1×. After fixation, HeLa cells were permeabilized and incubated in blocking buffer solution (5% w/v bovine serum albumin (BSA), 0.1% (v/v) Triton X-100 in PBS) for 1 hour at room temperature.

For PARP-1 detection, cells were incubated overnight at 4 °C with the primary antibody mouse anti-PARP1 (sc-8007; Santa Cruz Biotechnology), in blocking buffer (1/50 dilution), followed by several washing steps. Cells were then incubated with the secondary antibody Alexa488-conjugated anti-mouse (A28175; Thermofisher Scientific) in PBS (1/600 dilution), for 1 hour at room temperature, and washed with PBS.

Cells were stained with TO-PRO-3 Iodide (T3605; Thermofisher Scientific) (dilution 1:2000) in PBS and left incubating for 25 min at room temperature and subsequently were washed several times with ultrapure water.

### 2.5. Image processing and FRET calculation

All the following image operations were implemented on ImageJ.

For each measurement, the image of the phase lifetime *τ*(*x,y*) was obtained from the phasor images *g*(*x,y*) and *s*(*x,y*) using the formula:

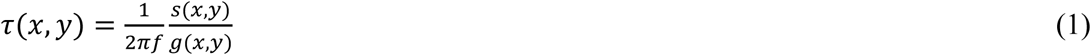

Where *f*=80 MHz. For donor-acceptor samples, the image of the FRET efficiency *E*(*x,y*) was obtained as:

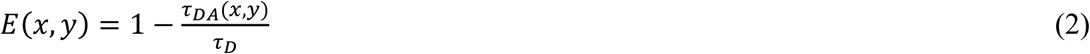

Where *τ*_DA_(*x,y*) is the phase lifetime image of a donor-acceptor sample and *τ*_D_ represents the phase lifetime of the unquenched donor. The lifetime of the unquenched donor was determined, in each experiment, from lifetime images of a donor only sample prepared in the same conditions of the donor-acceptor sample. The value *τ*_D_ was set as the average value obtained from at least 3 different cells.

The FRET level *A*(*x,y*) was then obtained as:

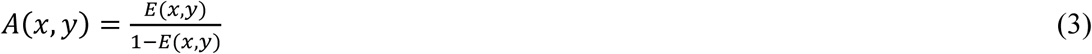

For donor-acceptor samples, the image of the acceptor-donor ratio was determined from the intensity images in the donor and acceptor channel. First, the contribution of the donor bleed-through was removed from the acceptor channel:

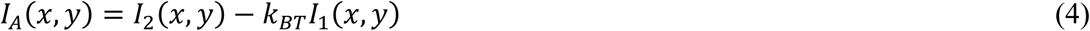

Where the constant *k*_BT_ was determined from the donor only sample as the average value of *I*_2_(*x,y*)/*I*_1_(*x,y*) from at least 3 different cells. The corrected intensity *I*_A_(*x,y*) represents all the fluorescence emission detected from Syto 13, including fluorescence resulting from direct excitation of Syto 13 at 405 nm and FRET signal. We will assume that the contribution of the FRET signal to *I*_A_(*x,y*) is negligible compared to the contribution due to direct excitation. The intensity in the donor channel was not affected by spectral cross-talk so we set *I*_D_(*x,y*)=*I*_1_(*x,y*). For the donor channel is *I*_D_(*x,y*)=*β*_D_*N*_D_(*x,y*)(*τ*_DA_(*x,y*)/*τ*_D_), where *β*_D_ is the brightness of the unquenched donor in the donor channel, *N*_D_(*x,y*) is the number of donor molecules at a given pixel and the factor *τ*_DA_/*τ*_D_ takes into account the decrease of quantum yield of the donor due to FRET [49]. For the acceptor channel is *I*_A_(*x,y*)=*β*_A_*N*_A_(*x,y*), where *β*_A_ is the brightness of the acceptor in the acceptor channel and *N*_A_(*x,y*) is the number of acceptor molecules at a given pixel. We calculated an image of the experimental acceptor-donor ratio as:

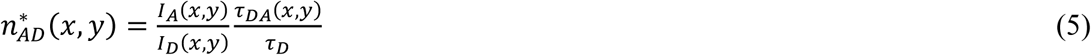

The quantity *n*\*(*x,y*) is proportional to the absolute value of the acceptor-donor ratio *N*_A_(*x,y*)/*N*_D_(*x,y*):

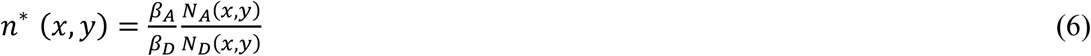

The value of *n*\*(*x,y*) was used to correct the measured FRET level *A*(*x,y*) for variations of the acceptor-donor ratio:

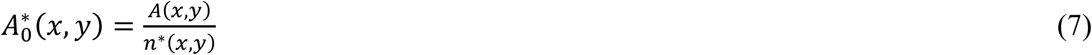

The quantity *A*_*0*_*(*x,y*) is proportional to the FRET level *A*_*0*_(*x,y*) corresponding to an acceptor-donor ratio of 1:

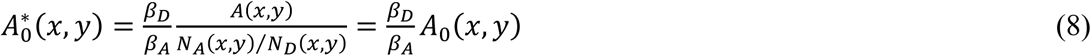

To estimate the value of the constant *β*_A_/*β*_D_, we first determined the ratio between the brightness of Fluorescein in the acceptor channel (*β*_2_) and the brightness of Alexa 405 in the acceptor channel (*β*_1_). This ratio was determined by measuring the fluorescence intensity *I*_1_ and *I*_2_ from distinct solutions of known concentration *C*_1_ and *C*_2_ of the two dyes under the same level of 405 nm excitation power.

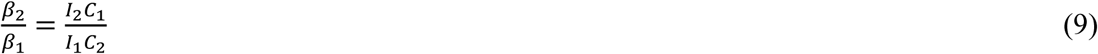

Then we estimated the value of *β*_A_/*β*_D_ as:

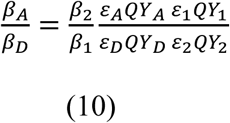

Where the values of quantum yield (QY) and extinction coefficient (ε) at 405 nm for each dye were estimated from reported literature and manufacturers datasheets (Supplementary Table 1).

### 2.6. Simulated data

The simulated data were generated in Matlab (Mathworks). We simulated a mixture of N_D_ donors undergoing FRET with a variable number N_A_ of acceptors. The lifetime of the unquenched donor was set to the value τ_0_=2.7 ns. The FRET efficiency of a donor interacting with a single acceptor was set to the value E_0_. The FRET efficiency of a donor interacting with an integer number *n*_A_ acceptors was set to the value E_n_, where E_n_=1/(1+1/A_n_), A_n_=*n*_A_A_0_ and A_0_=E_0_/(1−E_0_) [41]. For any given value of N_A_/N_D_, the mixture was set in the following way: for *n*_A_−1<N_A_/N_D_<*n*_A_, a value of efficiency E_n_ was assigned to a fraction of donors equal to N_A_/N_D_−*n*_A_, whereas a value of efficiency E_n−1_ was assigned to all the other donors. The temporal decay corresponding to this mixture was then analyzed in frequency domain via a Fast Fourier Transform algorithm. The phase value φ corresponding at the frequency *f*=80 MHz was used to calculate a value of phase lifetime τ_φ_=tanφ/(2π*f*). Finally, the efficiency E was calculated as E=1−τ_φ_/τ_0_ and the value of A was calculated as A=E/(1−E).

## 3. Results

### 3.1. A FRET assay corrected for variations in the acceptor-donor ratio

To measure chromatin compaction at the nanoscale level, in live cells, we explored the possibility of using the FRET occurring between nucleic acid binding dyes with overlapping emission/excitation spectra. An example of such a FRET pair is represented by Hoechst 33342 and Syto 13 (Fig.S1). Hoechst 33342 is a bisbenzimidazole dye binding to the minor groove of DNA with preferential AT-sequence specificity. Syto 13 labels DNA, both in the nucleus and mitochondria, and RNA in the cytoplasm and nucleoli. Consequently, the Syto dyes do not act exclusively as nuclear stains in live cells [50]. We assume that a higher nanoscale chromatin compaction corresponds, on average, to a shorter distance between the fluorophores and thus an increasing FRET efficiency (Fig.1). This is the rationale for using FRET as a quantitative measure of nanoscale chromatin compaction. On the other hand, an increase of the acceptor-to-donor abundance could produce by itself an increasing FRET efficiency, not necessarily related to the nanoscale chromatin compaction level (Fig.1). For this reason, it is important to extract a value of FRET efficiency related only to the average distance between the fluorophores but not to their relative abundance.

**Figure 1.**
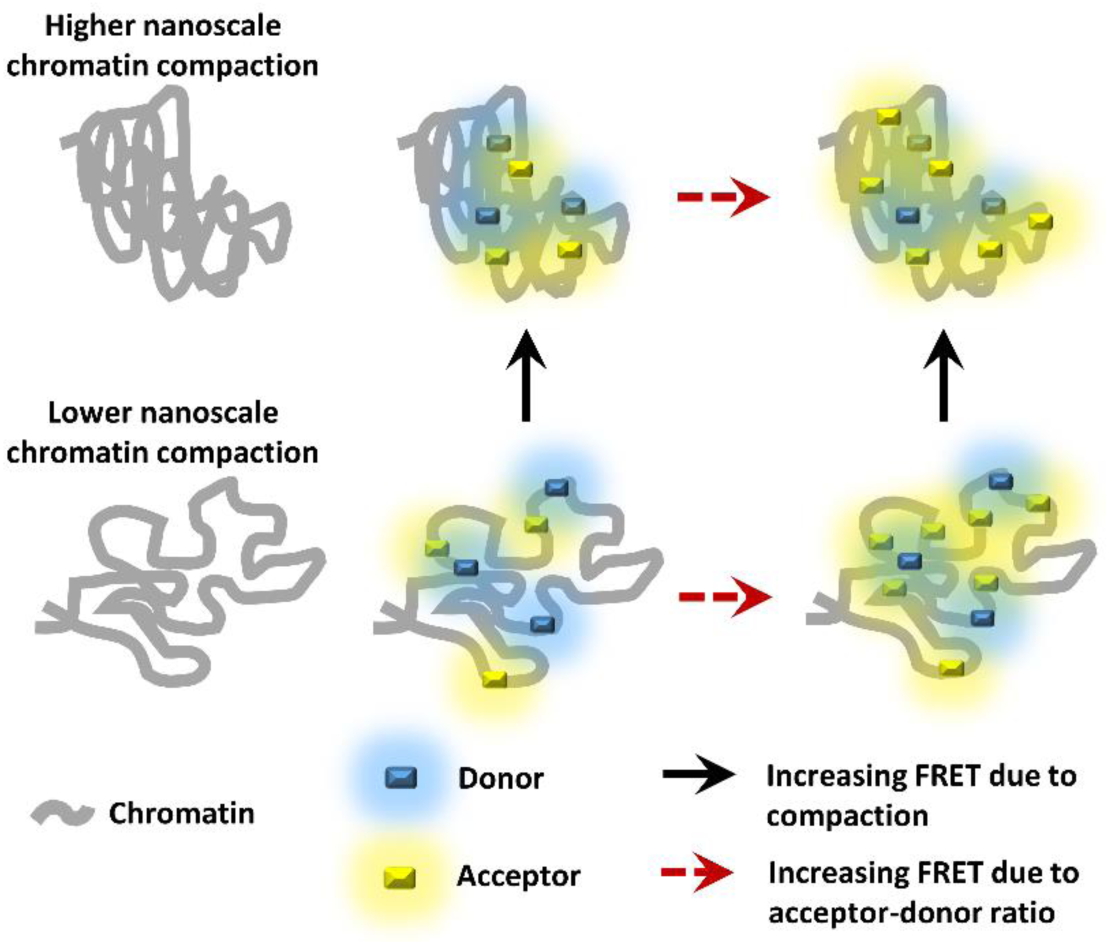
Schematic representation of the FRET assay for chromatin nanoscale compaction. Donor (cyan) and acceptor (yellow) molecules are bound to nuclear chromatin (grey). Variations in the measured FRET level can be due both to changes in chromatin density (black solid arrow) and/or to changes in relative acceptor-to-donor abundance (red dashed arrow).

To this aim, we established a protocol for measuring FRET at each pixel of an image and normalizing the FRET value to the relative acceptor-to-donor abundance (Fig.2). We measured FRET between Hoechst 33342 and Syto 13 by monitoring the decrease of the donor lifetime by frequency-domain FLIM. When FRET occurs, the donor (Hoechst 33342) lifetime is reduced due to energy transfer to the acceptor fluorophore (Syto 13). One advantage of FLIM is that it provides a value of FRET efficiency without the corrections (e.g. cross-talk between channels, estimation of the relative concentration of the fluorophores) required by intensity-based methods [51]. The FRET-induced decrease in the donor lifetime was detected by frequency-domain FLIM as a decrease in the value of phase measured at a given frequency (Fig. 2a). We observed that a higher acceptor-to-donor abundance corresponded to a stronger decrease of the donor lifetime (Fig. 2b-e). For instance, in the representative images shown in Fig.2, the average lifetime of Hoechst 33342 changed from τ_D_ ∼ 2.7 ns (donor only) to τ_DA_ ∼1.9 ns (Fig.2d) and τ_DA_ ∼ 1.7 ns (Fig.2e) respectively, due to FRET with Syto 13, resulting in different levels of the FRET efficiency *E*, defined as *E*=1−τ_DA_/τ_D_ (Fig.2i,j). These variations of FRET efficiency were related to variations in the relative acceptor-to-donor abundance naturally occurring on a given specimen (Fig.2d,e). To examine the dependence of the FRET level upon the relative acceptor-to-donor abundance is convenient to use the quantity *A*=*E*/(1-*E*) [41]. Simulated data showed indeed that the FRET level *A* is approximately linearly dependent upon the relative acceptor-to-donor abundance N_A_/N_D_, whereas the FRET efficiency *E* is not (Fig.2f). The slope of the *A* vs N_A_/N_D_ plot is a quantity dependent only from the acceptor-distance (Fig.2f, Fig.S2). The simulations also indicated that very different combinations of values of the parameters *E*_0_ (namely the efficiency of a donor coupled to a single acceptor) and N_A_/N_D_ can generate identical values of FRET level *A*. In other words, to determine the absolute value of *E*_0_ from the measured FRET level *A*, one must know the absolute value of the ratio N_A_/N_D_.

**Figure 2.**
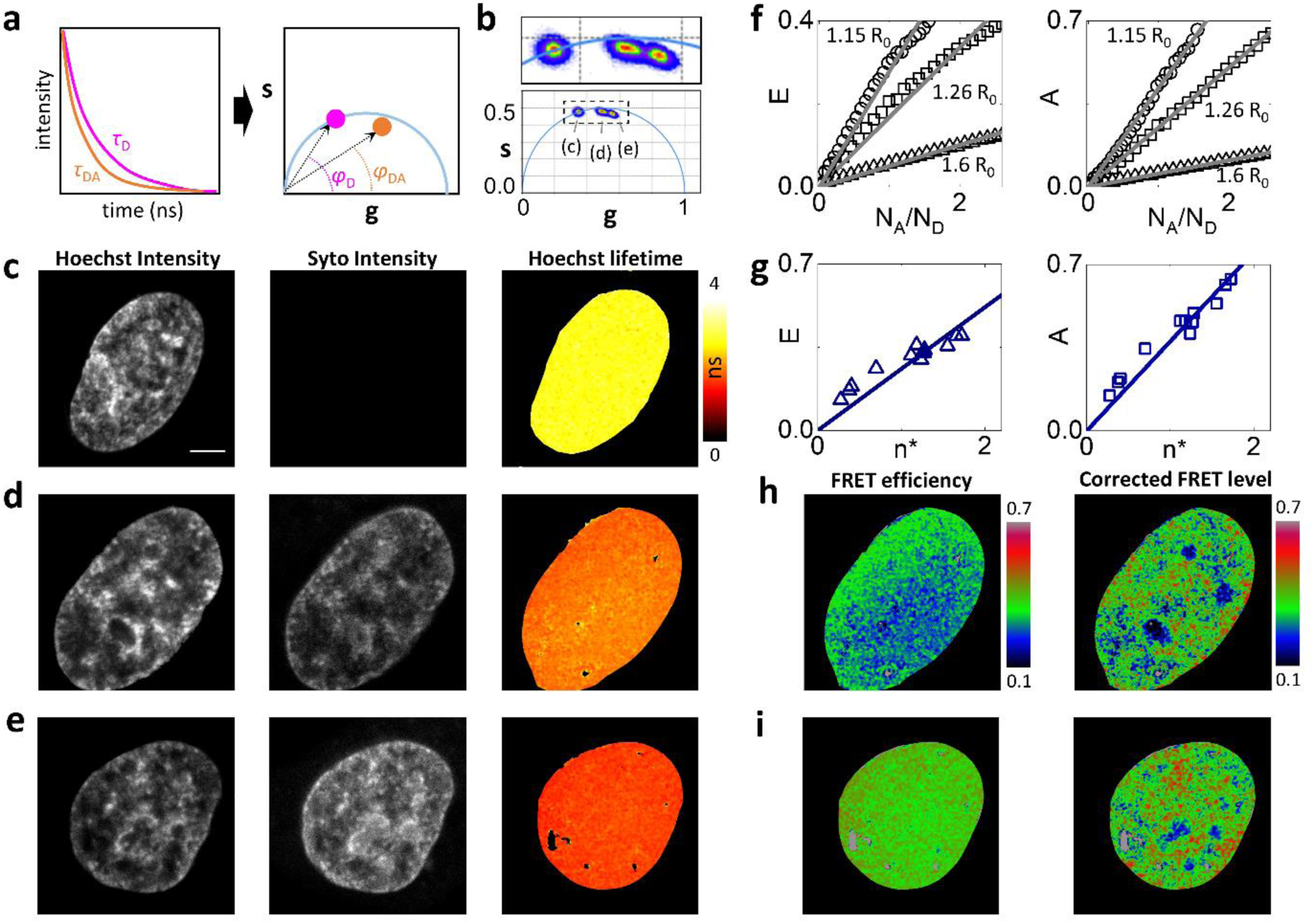
FRET assay corrected for the acceptor-donor ratio. (a) Schematic representation of frequency domain analysis of FLIM-FRET data. The FRET-induced decrease in the donor lifetime from τ_D_ to τ_DA_ is detected as a decrease in the value of phase (from φ_D_ to φ_DA_) measured at a given frequency. (b) Phasor analysis of FLIM-FRET of Hoechst 33342 and Syto 13 in live HeLa cells. The three clusters correspond to the representative samples reported in (c-e). (c-e) Representative images of a donor only sample (c) and of two donor-acceptor samples with different level of acceptor-donor ratio (d,e). Shown are the intensity in the donor channel, the intensity in the acceptor channel and the lifetime of the donor. (f) Plot of the FRET efficiency E and of the FRET level A versus the acceptor-donor ratio for simulated data. Numbers indicate the simulated acceptor-donor distance expressed in Forster radius (R_0_) units. Solid lines are linear fits of the data through the origin. (g) Experimental values of E and A versus n*, a quantity proportional to the acceptor-donor ratio. Solid lines are linear fits of the data through the origin. (h,i) FRET level before and after correction for the samples reported in (d,e). Shown are the map of the FRET efficiency E and the map of the corrected FRET level A_0_*. Scale bar: 5µm.

In our experimental data, we quantified the relative acceptor-to-donor abundance by a quantity proportional to the relative concentration of the two fluorophores, *n*\*= (*I*_A_/*I*_D_)(*τ*_DA_/*τ*_D_) ∝ *N*_A_/*N*_D_ (see Materials and Methods), where *I*_A_ and *I*_D_ are the intensity in the donor and acceptor channel respectively, and the factor τ_DA_/τ_D_ takes into account the decrease of quantum yield of the donor due to FRET [49]. The experimental dependence of the FRET efficiency *E* and the FRET level *A* versus the experimental acceptor-to-donor ratio n* are reported in Fig.2g. Finally, we used the quantity n* to normalize the FRET value *A* to the relative acceptor-to-donor ratio and generate a corrected value of FRET level A_0_*=*A*/n* at each pixel of an image (Fig.2h,i).

### 3.2. Regions of different chromatin density exhibit different nanoscale chromatin compaction

To validate the method, we first compared the values of FRET measured on chromatin regions of different DNA density. Regions of low and high density were identified, on each cell, based on the fluorescence intensity of Hoechst 33342 (Fig.3a,b). The regions corresponding to higher Hoechst 33342 signal typically included the peripheral and perinucleolar heterochromatin, because of the higher DNA concentration of these regions and also because of a preferential affinity of the dye towards AT-rich sequences [52]. The average FRET level A measured in the two regions, for each cell, was reported as a function of the relative acceptor-to-donor abundance n* (Fig.3c,e). The different slope of the two sets of data indicated that the regions with high Hoechst signal (the heterochromatin regions) have a higher level of nanoscale compaction compared to the regions with low Hoechst signal (the euchromatin regions). Similarly, the maps of corrected FRET level A_0_* (Fig.3d,f) showed that nanoscale compaction was higher in heterochromatin (A_0_*=0.42 ± 0.01, mean ±s.e.m., n = 10 cells) than in euchromatin (A_0_*=0.35 ± 0.01, n = 10 cells) (P<0.001, paired t-test, n = 10 cells). To test the sensitivity to alterations of the higher-order chromatin architecture we applied the approach used by Albiez et al.[46]. It consists in a modulation of chromatin compaction in living cells from normally condensed chromatin to hypercondensed chromatin by an increase of the osmolarity of the culture medium from 290 mOsm (standard osmolarity of normal growth medium) to 570 mOsm. This procedure resulted in an increase of chromatin condensation (Fig.3g,h). HeLa cells showed dense chromatin regions throughout the nucleus followed by a reduction in nuclear size. The average FRET level A was reported as a function of the relative acceptor-to-donor abundance n* (Fig.3i). The larger value of slope indicated that treatment with hyperosmolar solution had induced an increase in the nanoscale compaction as measured by FRET. Similarly, the maps of corrected FRET level A_0_* showed that nanoscale compaction was higher (P<0.001, t-test) in hypercondensed chromatin (A_0_*=0.63 ± 0.01, mean ±s.e.m., n = 11 cells) compared to control nuclei (A_0_*=0.37 ± 0.01, mean ±s.e.m., n = 8 cells) (Fig.3j).

**Figure 3.**
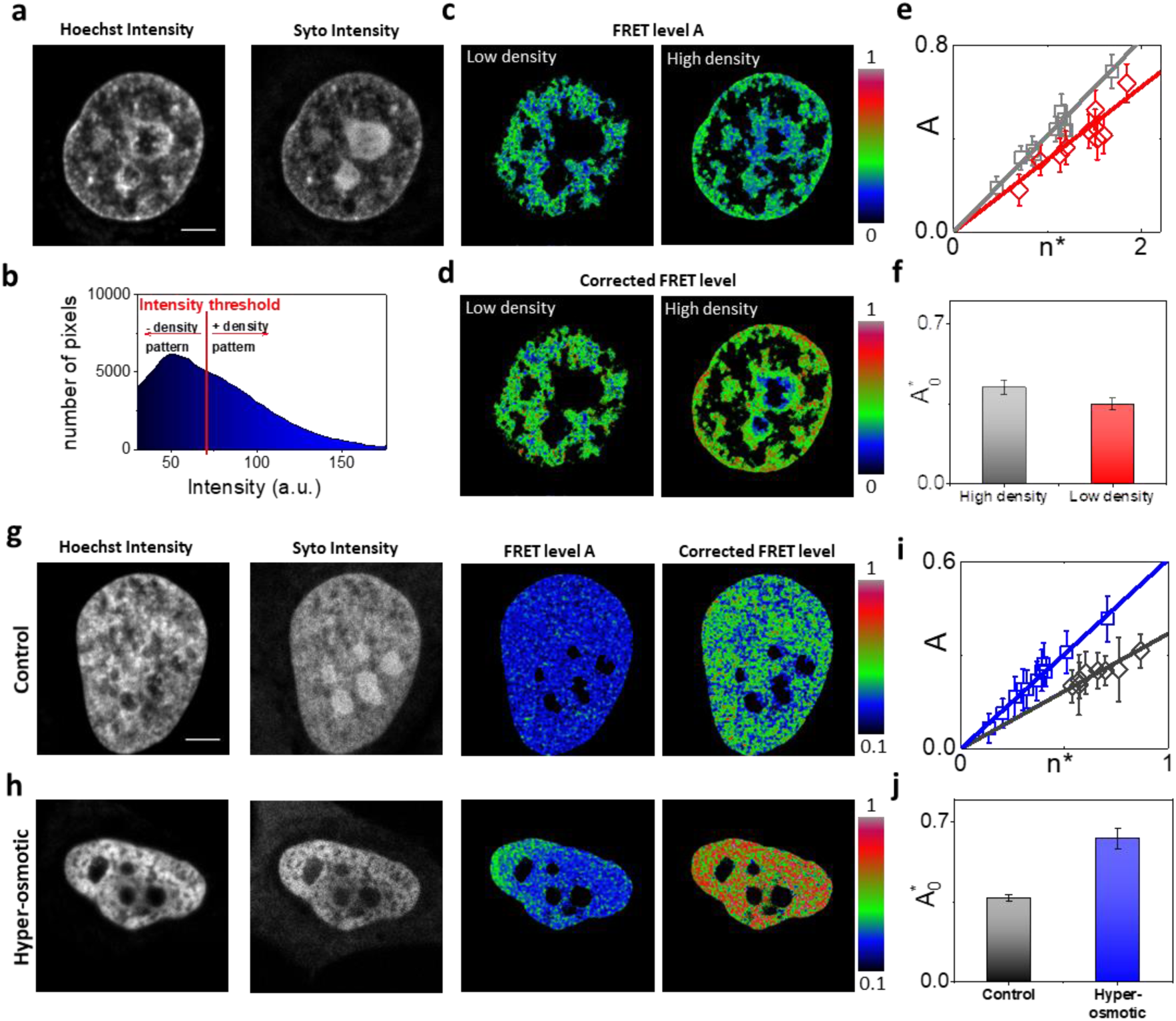
Corrected FRET assay shows different levels of nanoscale chromatin compaction. (a-f) FRET analysis of regions of different chromatin density in live HeLa cell nuclei. (a) Intensity images of the donor and acceptor channel. (b) A threshold in the Hoechst intensity histogram discriminates between high-density (heterochromatin) and low-density (euchromatin) regions. (c) Color maps of FRET level *A* in low density (left) and high density (right) regions. (d) Color maps of the corrected FRET level *A*_0_*. (e) Plot of *A* versus the relative acceptor-to-donor abundance *n*\*. Each experimental point is the mean ± s.d value of *A* calculated in a high (grey squares) or low (red diamonds) density region. The solid lines are linear fits of the data through the origin (high density: slope= 0.41; low density: slope= 0.31). (f) Comparison of mean value of A_0_* of heterochromatin (high density) and euchromatin (low density). Data are mean ± s.d. (n=10 cells) of the mean values of A_0_* calculated on each cell. (g-j) FRET analysis of hyper-osmotic nuclei. (g,h) Representative images of a control cell nucleus and a nucleus after hyper-osmolar treatment. Shown are the intensity in the donor and acceptor channel, the FRET level *A* and the corrected FRET level A_0_*. A_0_* color maps reveals and higher nanoscale compaction in hyper-osmotic nucleus with respect the control. (i) Plot of *A* versus the relative acceptor-to-donor abundance *n*\*. Each experimental point is the mean ± s.d value of *A* calculated in control (grey diamonds) or hyper-osmotic (blue squares) nuclei. The solid lines are linear fits of the data through the origin (control: slope= 0.37; hyper-osmotic: slope= 0.61) (j) Comparison of mean value of A_0_* of control and hyper-osmotic nuclei. Data are mean ± s.d. (control: n=8 cells; hyper-osmotic: n=11 cells) of the mean values of A_0_* calculated on each cell. Scale bar: 5µm.

### 3.3. Chromatin is decompacted at the nanoscale in response to DNA damage

Chromatin reorganization during DNA damage response (DDR) is a complex process. In particular, it has been previously reported that, following local induction of DNA damage, chromatin undergoes a rapid transient (<5 min) expansion followed by a slower compaction phase [53][54]. The rapid decondensation of chromatin in the early phase of DDR is a required step to allow the DNA-repair machinery to access the damaged region, thereby facilitating DNA damage repair [4]. Here, as an application of our FRET method, we tested if chromatin was locally decompacted, at the nanoscale, in response to induction of DNA damage.

To this aim, we generated DNA damage on a selected sub-region of the nuclei by 405nm-laser micro-irradiation, as reported previously [55]. To verify that 405nm-laser micro-irradiation generated local DNA damage in HeLa cell nuclei stained with Hoechst 33342, we monitored the expression of the DDR marker PARP-1 (poly-(ADP-ribose) polymerase1), a specific protein that rapidly accumulates at genome sites where single strand breaks (SSBs) or double strand breaks (DSBs) have occurred [3]. The expression of PARP-1 was monitored in cells fixed immediately after micro-irradiation and in live cells (Fig.4).

**Figure 4.**
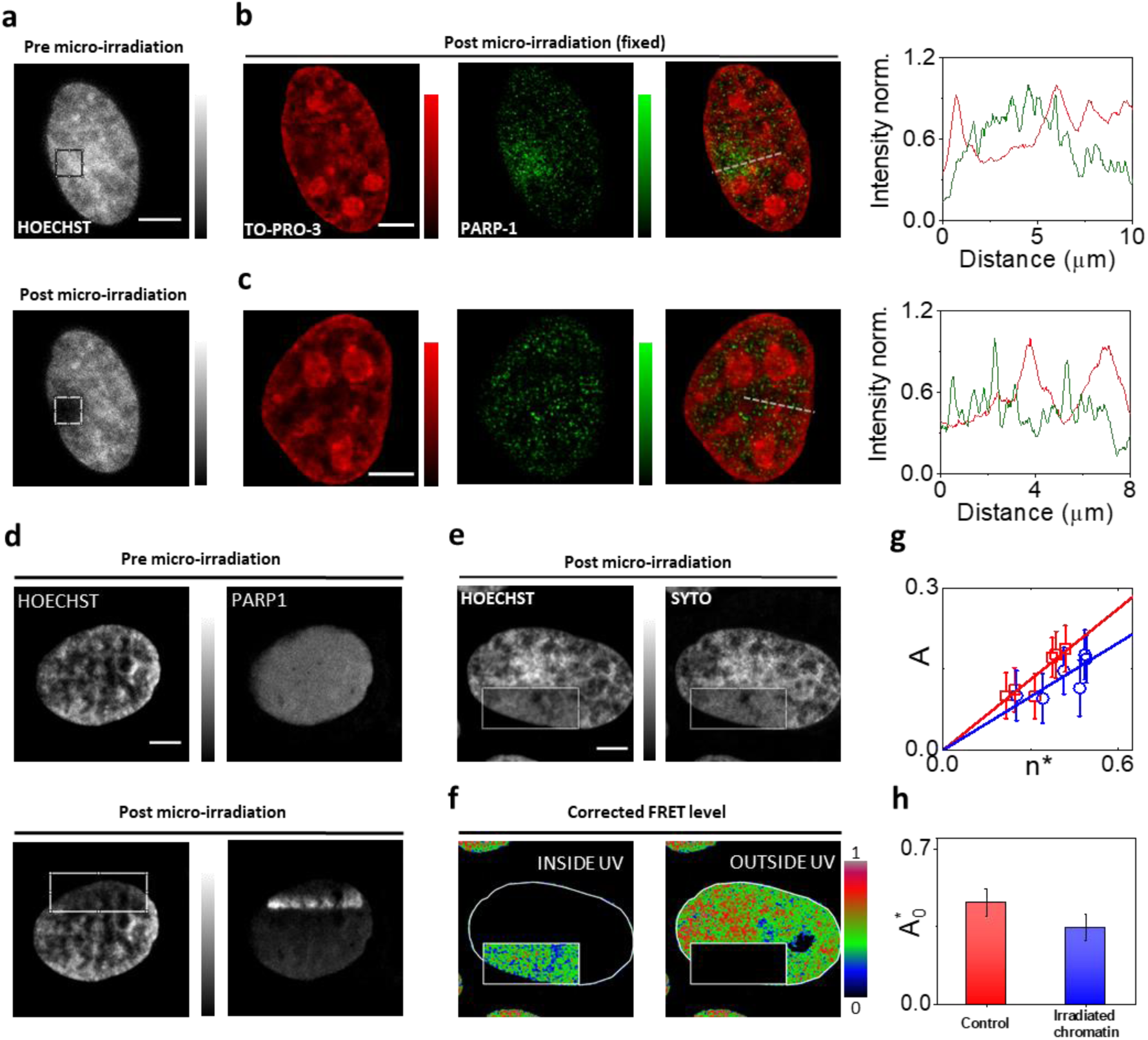
Corrected FRET assay reveals nanoscale decompaction of chromatin in response to DNA damage. (a) DNA damage is induced by UV-microirradiation in a region of interest (ROI) in a live HeLa cell nucleus labeled with Hoechst. (b,c) Confocal images showing DNA staining with TO-PRO-3 and immunodetection of PARP-1 in the same HeLa cell fixed in 4% PFA immediately after micro-irradiation (b) and in a control cell (c). Line profile shows the intensity signal distribution of TO-PRO-3 (red) and PARP-1 (green) in the irradiated region. (d) Representative images of a live HeLa cell nucleus stained with Hoechst and expressing PARP-1 chromobody tagged-RFP. Following local UV-microirradiation, there is accumulation of PARP-1 in the irradiated region. (e-h) FRET analysis in live HeLa cells stained with Hoechst and Syto 13 after local UV-microirradiation. (e) Intensity images of the donor and acceptor channel. (f) Color maps of the corrected FRET level (A_0_*) in selected masks representing the UV-irradiated ROI and outside ROI. (g) Plot of A versus the relative acceptor-to-donor abundance n*. Each experimental point is the mean ± s.d value of A calculated in UV-irradiated ROI (blue circles) or outside ROI (red squares) inside nuclei. The solid lines are linear fits of the data through the origin (UV-irradiated ROI: slope= 0.33; outside ROI: slope= 0.44) (h) Comparison of mean value of A_0_* of UV-irradiated and outside ROIs inside nuclei. Data are mean ± s.d. (n=6 cells) of the mean values of A_0_* calculated on each cell. Scale bar: 5µm.

In fixed cells, we observed accumulation of PARP-1 on the irradiated region (Fig. 4a-c). Post-fixation labeling with the DNA dye TO-PRO-3 [56] revealed, as expected, a local decondensation of DNA at the irradiation site (Fig. 4b). In live cells, we also observed photobleaching of Hoechst 33342 and accumulation of PARP-1 on the irradiated region (Fig.4d). The accumulation of PARP-1 was more prominent towards the center of the nucleus (Fig.4d and Supplementary Fig.3). One possible explanation for this effect is that more DNA damage is generated towards the center of the nucleus where the thickness is larger. In this respect, it is worth noting that, under one-photon excitation regime, the absorption of light and the subsequent generation of DNA damage is not limited to the focal plane but extended to the whole exposed volume [57].

We then measured the FRET level on HeLa cells stained with Hoechst 33342 and Syto 13 right after a region of the nucleus was exposed to laser micro-irradiation (Fig.4e-h). The FRET measurements were performed immediately after irradiation (within minutes), to focus only on the nanoscale rearrangement of chromatin occurring during the first expanding phase. The maps of corrected FRET level A_0_* showed that nanoscale compaction was lower in the exposed region of the nucleus (A_0_*=0.35 ± 0.02, mean ±s.e.m., n = 6 cells) compared to the non-exposed region (A_0_*=0.46 ± 0.02, mean ±s.e.m., n = 6 cells) (Fig.4f-h, P<0.005, paired t-test). The lower value of corrected FRET is an indication that chromatin is locally decompacted, at the nanoscale, in response to DNA damage induction, probably to promote the access of the DNA-repair machinery required for DNA damage repair, in keeping with reported models of chromatin organization [4].

## 4. Discussion

In this work, we have shown that a pair of DNA binding dyes, normally used as nuclear counterstains, can be used as a FRET system to map chromatin compaction within live cell nuclei. We have defined a successful strategy to distinguish the variations of FRET related to the donor-acceptor distance from the variations of FRET related to the acceptor-to-donor abundance. This strategy is based on a combination of both fluorescence lifetime and intensity measurements. The FRET level is quantified via the decrease of the donor lifetime, which is detected by FLIM, and then normalized to the relative acceptor-donor ratio, which is estimated from the intensity values in the acceptor and donor channels. To validate this strategy, we measured the FRET level in regions of high and low DNA density, as defined by the relative amount of Hoechst signal, and found that heterochromatin regions had a higher FRET level compared to the euchromatin regions. We also showed that nuclei of cells treated with a hyperosmolar medium had a higher FRET level compared to control nuclei. Finally, we applied our FRET method to monitor nanoscale reorganization of chromatin during response to DNA damage: we found that chromatin is locally decompacted, at the nanoscale, in response to DNA damage induction, probably to promote the access of the molecular machinery required for DNA damage repair.

These results show that, thanks to the normalization step, the reported FRET assay can be used to investigate chromatin organization in live cells, and is a valid alternative to the previously reported histone-based FRET systems. A major drawback, compared to the histone-based FRET assay, is that it is not straightforward to interpret our data in terms of the higher order organization of chromatin-DNA. Indeed, while the inter-nucleosome distance is a parameter directly connected to the chromatin higher order organization, here the precise spatial distribution of the two fluorophores on the DNA macromolecule is poorly defined. We can only make some simple assumptions on the distribution of the two binding dyes on DNA and attempt to estimate their average distance from the measured values of FRET efficiency.

FRET is a process sensitive to nanometer distances between fluorophores. However, the average FRET efficiency measured on a given pixel depends also on how many donors and acceptors are engaged in the FRET interaction. For instance, in protein-protein interactions it is common to describe FRET data in terms of a mixture of two species: a fraction of unquenched donors and a fraction of donors undergoing FRET with an acceptor [33]. Here, we cannot make any prior hypothesis on the stoichiometry of the FRET interaction. In order to convert the measured values of FRET efficiency into nanometer values, it is necessary to know the absolute value of the ratio N_A_/N_D_. The relationship between N_A_/N_D_ and our experimental parameter n* is given by N_A_/N_D_=(β_D_/β_A_)×n*, where β_A_ and β _D_ are the brightness of the acceptor and of the unquenched donor in the acceptor and donor channel, respectively. The simplest way to estimate the ratio β_D_/β_A_ would be to use a specimen with a 1:1 stoichiometry of the two fluorophores [58]. This is particularly challenging in our system since the brightness of a dye bound to DNA is much higher than that of the free dye and the effective concentration of dye bound to DNA can be very different from that of the staining solution. Using solutions of known concentration of organic dyes with similar emission spectra, we estimated indirectly that, in our system, β_D_/β_A_∼3.5, resulting in absolute values of acceptor-to-donor ratios N_A_/N_D_ ranging between ∼1 and ∼5 in our experiments. If this estimation is correct, we can then calculate the FRET value corresponding to a donor-acceptor pair as *A*_0_=*A*_0_*/(β_D_/β_A_). For the control samples, this value is in the order of *A*_0_∼0.1, corresponding to a FRET efficiency in the order of *E*_0_∼0.1 and average donor-acceptor distances in the order of ∼1.5 Forster radii. Assuming a Forster radius of ∼5 nm, this corresponds to ∼8 nm. According to recent electron microscopy observations, chromatin can be described a disordered chain with diameters between 5 and 24 nm, packed together at different concentration densities in interphase nuclei [59]. We can speculate that variations in the local density of this chain determine variations of the average acceptor-donor distance and thus variations in the detected FRET. In this framework, a variation of *A*_0_ from 0.1 to 0.12, like that observed between euchromatin and heterochromatin (Fig.3), would correspond to a variation of average acceptor-donor distance from ∼1.5 to only ∼1.4 Forster radii. This variation of distance can seem relatively small if compared with an estimated 2.6-fold difference of total DNA density between the two compartments [20]. However, this is not surprising considering the heterogeneity in the nanodomain size recently observed for DNA and nucleosome higher-order structures [16][20][59]. Our estimation of the average interaction distance might be inaccurate, as it does not rely on a robust calibration protocol. In this respect, we believe it would be interesting to perform a similar analysis on the histone-based FRET assays [36][37][60] where the use of fluorescent proteins would allow a more robust calibration with constructs of known stoichiometry [58] and a direct estimation of the average nanometer distance between labeled histones. Another interesting perspective is the combination of our FRET method with super-resolution microscopy. For instance, super-resolved FLIM-FRET measurements could be performed thanks to the recent integration of FLIM with Image Scanning Microscopy (ISM)[61], a technique similar to SIM that doesn’t require the use of special fluorophores. Finally, we believe that our method could be further improved with the use of novel fluorogenic dyes recently developed for a more efficient staining of chromatin [62]. At the same time, the compatibility of these new dyes with STED super-resolution microscopy opens the intriguing possibility of using the same fluorophore for imaging chromatin on different nanoscale windows: from the diffraction limit down to the tens of nanometers by STED and in the nanometer range by FRET.

## Supporting information

Additional supporting information may be found in the online version of this article at the JBP website. File containing Supporting Figures S1-S4 and Supplementary Table 1 (PDF).

## Authors contributions

LL and AD designed research. LL and SP performed experiments. LL and SP wrote software and analyzed data. The manuscript was written through contributions of all authors.

## Acknowledgements

LL was supported by Fondazione Cariplo and Associazione Italiana per la Ricerca sul Cancro (AIRC) through Trideo (Transforming Ideas in Oncological Research) Grant number 17215. The authors would like to thank Michele Oneto (IIT), Paolo Bianchini (IIT), Marco Scotto (IIT) and Ulas Coskun (ISS) for technical support.

## References

[1] T. Misteli, CSH Perspect. Biol., 2, a000794–a000794 (2010).

[2] T. Sexton, H. Schober, P. Fraser, and S.M. Gasser, Nat. Struct. Mol. Biol., 14, 1049 (2007).

[3] A.R. Chaudhuri and A. Nussenzweig, Nat. Rev., 18, 610–621 (2017).

[4] B.D. Price and A.D.D. Andrea, Cell, 152, 1344–1354 (2014).

[5] A. Seeber and S.M. Gasser, Curr. Opin. Genet. Dev., 43, 9–16 (2017).

[6] R. Imai, T. Nozaki, T. Tani, K. Kaizu, K. Hibino, S. Ide, S. Tamura, K. Takahashi, M. Shribak, and K. Maeshima, Mol. Biol. Cell, 28, 3349–3359 (2017).

[7] E. Fedorova and D. Zink, Biochim. Biophys. Acta, 1783, 2174–2184 (2008).

[8] S.I.S. Grewal and S.C.R. Elgin, Curr. Opin. Genet. Dev., 12, 178–187 (2002).

[9] S. W Hell and J. Wichmann, “Breaking the Diffraction Resolution Limit by Stimulated-Emission—Stimulated-Emission-Depletion Fluorescence Microscopy”, (1994).

[10] M.G.L. Gustafsson, J. Microsc., 198, 82–87 (2000).

[11] E. Betzig, G.H. Patterson, R. Sougrat, O.W. Lindwasser, S. Olenych, J.S. Bonifacino, M.W. Davidson, J. Lippincott-schwartz, and H.F. Hess, Science, 313, 1642–1646 (2006).

[12] M.J. Rust, M. Bates, and X. Zhuang, Nat. Methods, 3, 793 (2006).

[13] S. van de Linde, A. Löschberger, T. Klein, M. Heidbreder, S. Wolter, M. Heilemann, and M. Sauer, Nat. Protoc., 6, 991 (2011).

[14] B. Dong, L.M. Almassalha, Y. Stypula-cyrus, B.E. Urban, and J.E. Chandler, Proc. Natl. Acad. Sci., 113, 9716–9721 (2016).

[15] A.N. Boettiger, B. Bintu, J.R. Moffitt, S. Wang, B.J. Beliveau, G. Fudenberg, M. Imakaev, L.A. Mirny, C.T. Wu, and X. Zhuang, Nature, 529, 418–422 (2016).

[16] M.A. Ricci, C. Manzo, M. Lakadamyali, and M.P. Cosma, Cell, 160, 1145–1158 (2015).

[17] T. Nozaki, R. Imai, M. Tanbo, R. Nagashima, S. Tamura, and T. Tani, Mol. Cell, 67, 282–293.e7 (2017).

[18] L. Schermelleh, P.M. Carlton, S. Haase, L. Shao, P. Kner, B. Burke, M.C. Cardoso, D.A. Agard, G.L. Mats, H. Leonhardt, and J.W. Sedat, Science, 320, 1332–1336 (2010).

[19] M.J. Sarmento, M. Oneto, S. Pelicci, L. Pesce, L. Scipioni, M. Faretta, L. Furia, G.I. Dellino, P.G. Pelicci, P. Bianchini, A. Diaspro, and L. Lanzanò, Nat. Commun., 9, 1–11 (2018).

[20] K. Fang, X. Chen, X. Li, Y. Shen, J. Sun, D.M. Czajkowsky, and Z. Shao, ACS Nano, 12, 4909–4918 (2018).

[21] J. Xu, H. Ma, J. Jin, S. Uttam, R. Fu, Y. Huang, and Y. Liu, Cell Rep., 24, 873–882 (2018).

[22] T. Misteli, Science, 291, 843–847 (2001).

[23] L. Scipioni, M. Di Bona, G. Vicidomini, and L. Lanzanò, Commun. Biol., 1, 1–10 (2018).

[24] M. Baum, F. Erdel, M. Wachsmuth, and K. Rippe, Nat. Commun., 5, 1–12 (2014).

[25] N. Dross, C. Spriet, M. Zwerger, G. Müller, W. Waldeck, and J. Langowski, PLoS One, 4, e5041 (2009).

[26] A. Bancaud, S. Huet, N. Daigle, J. Mozziconacci, J. Beaudouin, and J. Ellenberg, EMBO J., 28, 3785–3798 (2009).

[27] M. Di Bona, M. Mancini, D. Mazza, G. Vicidominii, A. Diaspro, and L. Lanzanò, Biophys. J., 116, 1–13 (2019).

[28] C. Eggeling, C. Ringemann, R. Medda, G. Schwarzmann, K. Sandhoff, S. Polyakova, V.N. Belov, B. Hein, C. von Middendorff, A. Schönle, and S.W. Hell, Nature, 457, 1159 (2008).

[29] L. Lanzanò, L. Scipioni, M. Di Bona, P. Bianchini, R. Bizzarri, F. Cardarelli, A. Diaspro, and G. Vicidomini, Nat. Commun., 8, 65 (2017).

[30] S.T. Spagnol and K.N. Dahl, PLoS One, 11, 1–19 (2016).

[31] B. Banerjee, D. Bhattacharya, and G. V Shivashankar, Biophys. J., 91, 2297–2303 (2006).

[32] G. Bunt and F.S. Wouters, Biophys Rev, 9, 119–129 (2017).

[33] H. Giral, D.A. Cranston, L. Lanzano, Y. Caldas, E. Sutherland, J. Rachelson, E. Dobrinskikh, E.J. Weinman, R.B. Doctor, E. Gratton, and M. Levi, J. Biol. Chem., 287, 35047–35056 (2012).

[34] Y. Sun, H. Wallrabe, S.-A. Seo, and A. Periasamy, ChemPhysChem, 12, 462–474 (2011).

[35] H. Giral, L. Lanzano, Y. Caldas, J. Blaine, J.W. Verlander, T. Lei, E. Gratton, and M. Levi, J. Biol. Chem., 286, 15032–15042 (2011).

[36] D. Llères, J. James, S. Swift, D.G. Norman, and A.I. Lamond, J. Cell Biol., 187, 481–96 (2009).

[37] D. Llères, A.P. Bailly, A. Perrin, D.G. Norman, D.P. Xirodimas, and R. Feil, Cell Rep., 18, 1791–1803 (2017).

[38] M.A. Digman, V.R. Caiolfa, M. Zamai, and E. Gratton, Biophys. J., 94, L14–L16 (2008).

[39] J. Lou, L. Scipioni, B.K. Wright, T.K. Bartolec, J. Zhang, V.P. Masamsetti, K. Gaus, E. Gratton, A.J. Cesare, and E. Hinde, Proc. Natl. Acad. Sci., 116, 7323 LP–7332 (2019).

[40] K.H. Rainey and G.H. Patterson, Proc. Natl. Acad. Sci., 116, 864–873 (2019).

[41] Á.I. Fábián, T. Rente, J. SzölloSi, L. Matyus, and A. Jenei, ChemPhysChem, 11, 3713–3721 (2010).

[42] P. V Nazarov, R.B.M. Koehorst, W.L. Vos, V. V Apanasovich, and M.A. Hemminga, Biophys. J., 91, 454–466 (2006).

[43] S. V Koushik, P.S. Blank, and S.S. Vogel, PLoS One, 4, 1–12 (2009).

[44] C. Berney and G. Danuser, Biophys. J., 84, 3992–4010 (2003).

[45] A. Zeug, A. Woehler, E. Neher, and E.G. Ponimaskin, Biophys. J., 103, 1821–1827 (2012).

[46] H. Albiez, M. Cremer, C. Tiberi, L. Vecchio, L. Schermelleh, S. Dittrich, K. Küpper, B. Joffe, T. Thormeyer, J. Von Hase, S. Yang, K. Rohr, H. Leonhardt, I. Solovei, C. Cremer, S. Fakan, and T. Cremer, Chromosom. Res., 14, 707–733 (2006).

[47] A. Esposito, H.C. Gerritsen, and F.S. Wouters, Biophys. J., 89, 4286–4299 (2005).

[48] J. Schindelin, I. Arganda-Carreras, E. Frise, V. Kaynig, M. Longair, T. Pietzsch, S. Preibisch, C. Rueden, S. Saalfeld, B. Schmid, J.-Y. Tinevez, D.J. White, V. Hartenstein, K. Eliceiri, P. Tomancak, and A. Cardona, Nat. Methods, 9, 676 (2012).

[49] G. Vámosi, N. Baudendistel, C.-W. von der Lieth, N. Szalóki, G. Mocsár, G. Müller, P. Brázda, W. Waldeck, S. Damjanovich, J. Langowski, and K. Tóth, Biophys. J., 94, 2859–2868 (2008).

[50] M.A.M.J. Van Zandvoort, C.J. De Grauw, H.C. Gerritsen, J.L.V. Broers, M.G.A. Oude Egbrink, F.C.S. Ramaekers, and D.W. Slaaf, Cytometry, 47, 226–235 (2002).

[51] J.A. Broussard, B. Rappaz, D.J. Webb, and C.M. Brown, Nat. Protoc., 8, 265–281 (2013).

[52] G. Mascetti, L. Vergani, A. Diaspro, S. Carrara, G. Radicchi, and C. Nicolini, Cytometry, 23, 110–119 (1996).

[53] M.J. Kruhlak, A. Celeste, G. Dellaire, O. Fernandez-capetillo, W.G. Müller, J.G. Mcnally, D.P. Bazett-jones, and A. Nussenzweig, J. Cell Biol., 172, 823–834 (2006).

[54] R.C. Burgess, B. Burman, M.J. Kruhlak, T. Misteli, R.C. Burgess, B. Burman, M.J. Kruhlak, and T. Misteli, CellReports, 9, 1703–1717 (2014).

[55] C. Dinant, M. De Jager, J. Essers, W.A. Van Cappellen, R. Kanaar, A.B. Houtsmuller, and W. Vermeulen, J. Cell Sci., 120, 2731–2740 (2007).

[56] R.M. Martin, H. Leonhardt, and M.C. Cardoso, Cytometry, 67A, 45–52 (2005).

[57] A. Diaspro, G. Chirico, and M. Collini, Q. Rev. Biophys., 38, 97–166 (2005).

[58] M. Renz, B.R. Daniels, G. Vámosi, I.M. Arias, and J. Lippincott-schwartz, Proc. Natl. Acad. Sci., 1–9 (2012).

[59] H.D. Ou, S. Phan, T.J. Deerinck, A. Thor, M.H. Ellisman, and C.C.O. Shea, Science, 357, 1–13 (2017).

[60] J. Lou, L. Scipioni, B.K. Wright, T.K. Bartolec, J. Zhang, V.P. Masamsetti, K. Gaus, E. Gratton, A.J. Cesare, and E. Hinde, bioRxiv, 419523 (2018).

[61] M. Castello, G. Tortarolo, M. Buttafava, T. Deguchi, F. Villa, S. Koho, L. Pesce, M. Oneto, S. Pelicci, L. Lanzanó, P. Bianchini, C.J.R. Sheppard, A. Diaspro, A. Tosi, and G. Vicidomini, Nat. Methods, 16, 175–178 (2019).

[62] J. Bucevičius, J. Keller-Findeisen, T. Gilat, S.W. Hell, and G. Lukinavičius, Chem. Sci., (2019).

