## Supplemental Info for "Chromatin nanoscale compaction in live cells visualized by acceptor-donor ratio corrected FRET between DNA dyes"

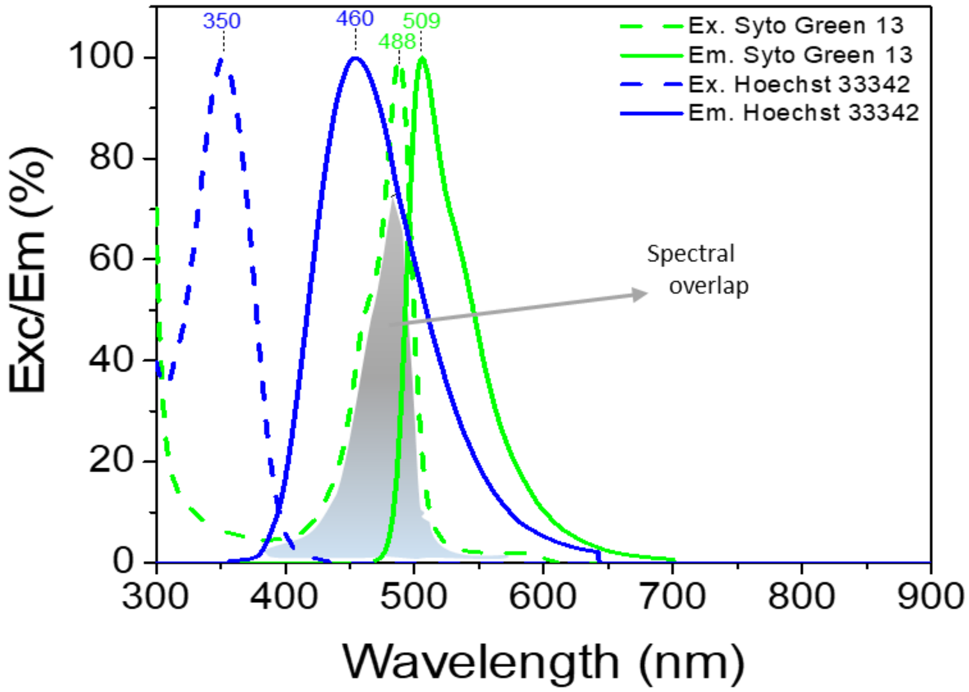


**Fig.S1**. Excitation and emission spectra of Hoechst 33342 and Syto Green 13 (Source: thermofisher.com). The shaded region represents the spectral overlap between Hoechst 33342 emission and Syto Green 13 absorption.

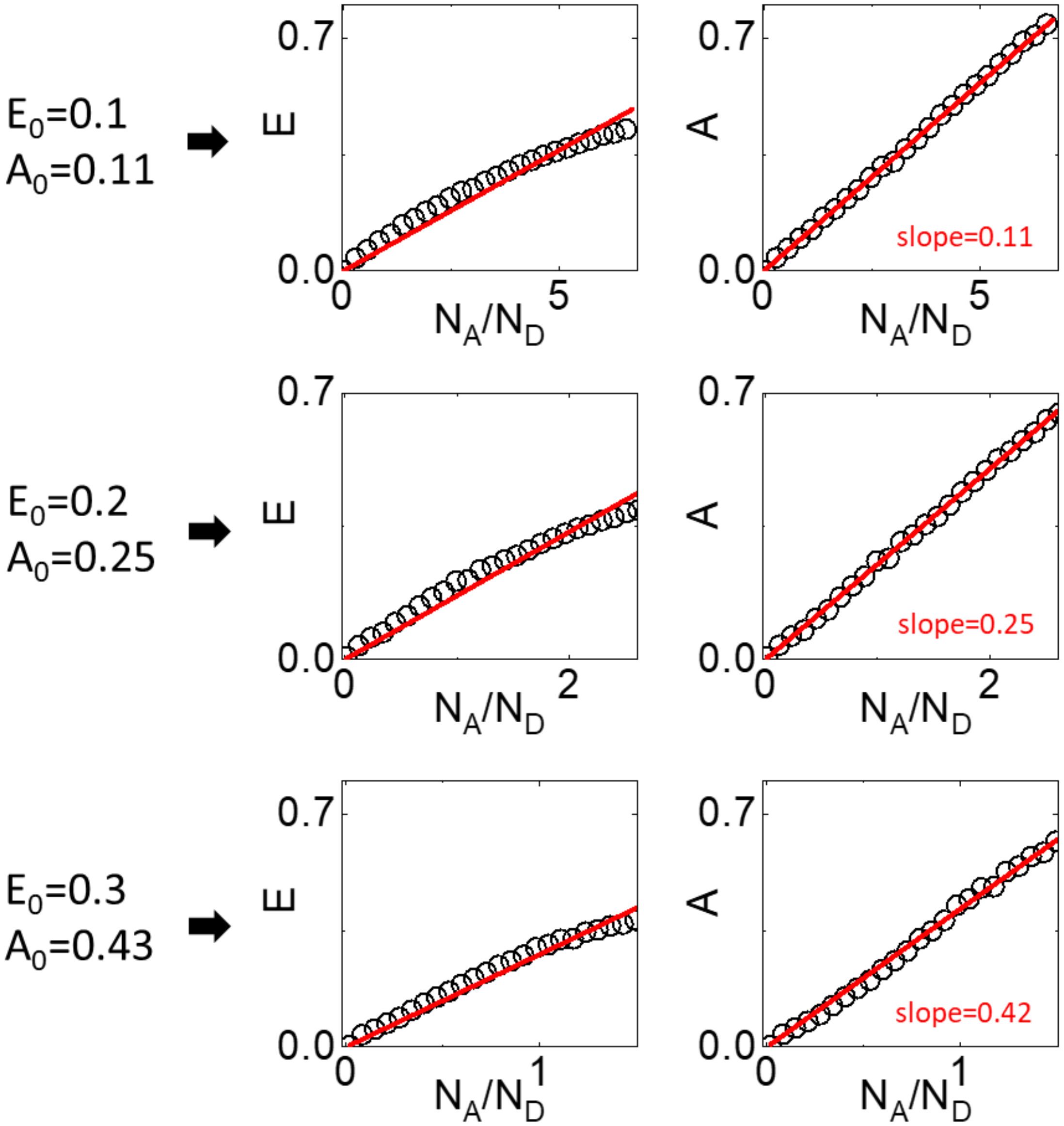


**Fig.S2**. Calculation of FRET efficiency E and FRET level A on simulated data. Data represent the FRET measured from a mixture of $N_D$ donors undergoing FRET with a variable number $N_A$ of acceptors (see details on Main Text). Each simulation has been obtained by fixing the value of $E_0$, corresponding to the FRET efficiency of a donor interacting with a single acceptor, and by varying the acceptor donor ratio $N_A/N_D$. The solid red lines are linear fit of the data through the origin.

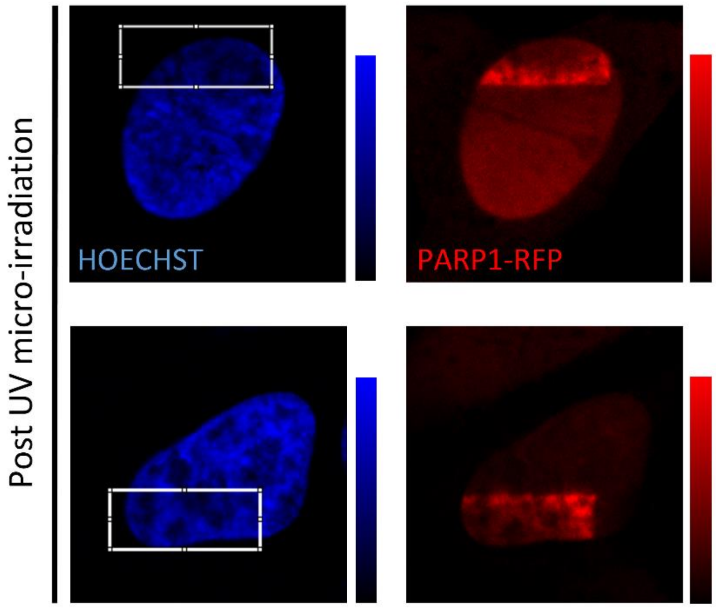


**Fig.S3**. Localization of PARP-1 in response to laser-induced DNA damage. Representative confocal images of live HeLa cell nuclei stained with Hoechst and expressing PARP-1 chromobody tagged-RFP, after exposure of the selected ROI to UV-microirradiation. The images show accumulation of PARP-1 in the irradiated region.

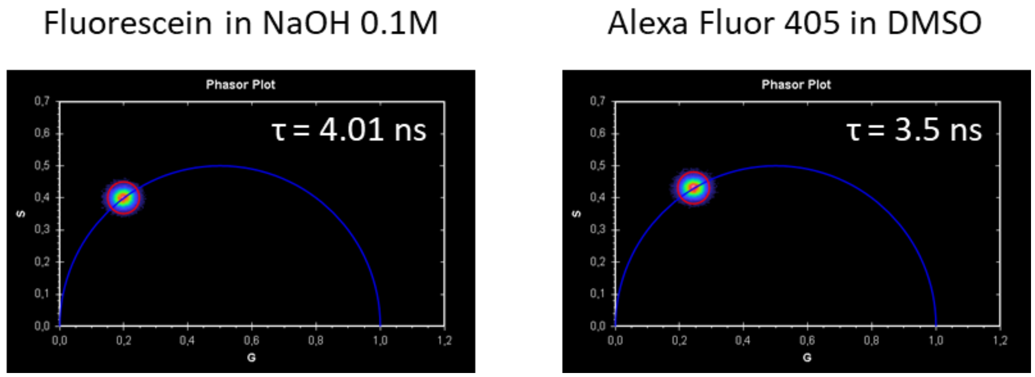


**Fig.S4**. Determination of the fluorescence lifetime of Alexa 405 in DMSO. The lifetime has been measured using 405 nm excitation modulated at 80 MHz. Shown is the phasor plot of a solution of Fluorescein in NaOH 0.1M used for calibration (left) and the phasor plot of a solution of Alexa 405 in DMSO (right).

**Supplementary Table 1**. Parameters related to the dyes Alexa Fluor 405 in DMSO, fluorescein in 0.1M NaOH, Hoechst 33342 bound to DNA, Syto 13 bound to DNA. Reported are the quantum yield (QY), the extinction coefficient at the absorption peak ($\varepsilon$), the percentage of absorption at $\lambda$=405 nm relative to the absorption peak and the extinction coefficient at $\lambda$=405 nm ($\varepsilon_{405}$).

| **DYE** | **Q.Y.** | **$\varepsilon$** | **% abs (405 nm)** | **$\varepsilon_{405}$** |
|---|---|---|---|---|
| Alexa Fluor 405 | 0.54 | 35000 $M^{-1}cm^{-1}$ | 86% | 30100 $M^{-1}cm^{-1}$ |
| Fluorescein | 0.93 | 76900 $M^{-1}cm^{-1}$ | 2% | 1540 $M^{-1}cm^{-1}$ |
| Hoechst 33342 | 0.38 | 47000 $M^{-1}cm^{-1}$ | 3.5% | 1645 $M^{-1}cm^{-1}$ |
| Syto 13 | 0.4 | 50000 $M^{-1}cm^{-1}$ | 5% | 2500 $M^{-1}cm^{-1}$ |
